# Post-copulatory behavior of olive baboons (*Papio anubis*) infected by *Treponema pallidum*

**DOI:** 10.1101/2020.02.23.961326

**Authors:** Filipa M. D. Paciência, Idrissa S. Chuma, Iddi F. Lipende, Sascha Knauf, Dietmar Zinner

## Abstract

In nonhuman primates pathogens are known to exert a profound and pervasive cost on various aspects of their sociality and reproduction. In olive baboons (*Papio anubis*) at Lake Manyara National Park, genital skin ulcers caused by *Treponema pallidum* subsp. *pertenue* lead to mating avoidance in females and altered mating patterns at a pre-copulatory and copulatory level. Beyond this level, sexual behavior comprises also post-copulatory interactions among the sexual partners. To investigate whether the presence of genital skin ulcers has an impact at the post-copulatory level, we analyzed 517 copulation events of 32 cycling females and 29 males. The occurrence of post-copulatory behaviors (i.e., copulation calls, darting [female rapid withdraw from the male] and post-copulatory grooming) was not altered by the presence of genital skin ulcerations. Similarly to other baboon populations, females of our group were more likely to utter copulation calls after ejaculatory copulation. The likelihood of darting was higher after ejaculatory copulations and with the presence of copulation calls. Post-copulatory grooming was rarely observed but when it occurred, males groomed females for longer periods when females uttered copulation calls during, or preceding mating. Our results indicate that despite the presence of conspicuous genital skin ulcers, the post-copulatory behavior was not affected by the genital health status of the dyad. This suggests that infection cues play a major role before and during mating but do not affect post-copulatory behavior.

## 1 INTRODUCTION

Mating is intrinsically associated with sexual selection, which occurs through competition over mates or mate choice (Darwin, 1871; Anderson, 1994). Mate choice might confer both direct and indirect fitness benefits to the choosy individual. Such benefits might be a higher level of parental care or the accumulation of “good genes” in the offspring (Andersson, 1994, Kokko et al., 2003). One important fitness criterion in the mating context is the health status of the potential partner. Thus, individuals should choose healthy partners, since mating with a sick individual may not only have negative effects on the offspring (i.e. a poor health status can be an indication of a poor immune system which would then passed on to the offspring), but also on the health of the choosy individual itself, if the disease can be transmitted (Hillgarth, 1996; Martinez-Padilla et al., 2012). The latter becomes particularly obvious if the disease is sexually transmitted. A poor health status can alter both the individual’s attractiveness as a sexual partner, and its competitive ability and performance in the mating context (Beltran-Bech et al., 2014). This is particularly true when courtship and mating are energetically demanding phases (Key & Ross 1999).

In some primate species, males follow females for hours and days, maintain close proximity, increase their grooming bouts and try to monopolize mating (e.g., Smuts, 1987; Dixson, 2013). Such an investment comes along with time and energy costs, which diseased males might not be able to cover. Likewise, females might not engage with males during copulation, e.g., not showing proceptive and receptive behavior, not permit subsequent matings or not showing interest in post-copulatory grooming.

In baboons, copulations are usually defined by male mounting with intromission and pelvic thrusts upon the female, which can culminate with ejaculation (Saayman, 1970, Ransom, 1981). Yet, the number of mounts and pelvic thrusts per mount may vary. An ejaculatory mount is usually identified by an ejaculation pause, where the male remains rigid upon the female for a few seconds (Saayman, 1970). During or after copulation, females might utter copulation calls which are typically low-frequency rhythmic vocalizations (Bouquet et al., 2018). In addition, female baboons often exhibit a characteristic post-coital sprint over several meters away from the male, a post-copulatory withdraw-behavior that is known as ‘darting’ (Hall & DeVore 1965; Saayman, 1970; Ransom, 1981; Smuts, 1985; Bercovitch, 1995; Collins, 1981; O’Connell & Cowlishaw 1995). Finally, pairs may engage in post-copulatory grooming (PCG), which can be initiated either by the male or the female (Saayman, 1970).

At Lake Manyara National Park (LMNP), olive baboons (*Papio anubis*) are infected with a putative sexually transmitted infection caused by the bacterium *Treponema pallidum* subsp. *pertenue* (*TPE*, Knauf et al., 2012, 2018; Harper et al., 2012; Chuma et al., 2016). Clinical symptoms are characterized by genital skin ulcers (in the following referred to as genital ulcers), a moderate to severe ulceration of the anogenital skin in both males and females (Figure 1). Progressive scarification of the tissue can lead to a permanently open state of the vagina and anus in females; while in males it cause phimosis or loss of the corpus penis (Knauf et al., 2012). At LMNP, genital ulcers in baboons have been linked with mating avoidance by females and altered copulatory patterns by males, i.e., ulcerated individuals exhibit fewer pelvic thrusts (Paciência et al., 2019). Since *TPE* infection has been associated with pre-copulatory mate choice, we aimed to investigate whether the post-copulatory behavior is altered by the genital health status (i.e. presence of genital ulcers) of the mating pairs.

**FIGURE 1.**
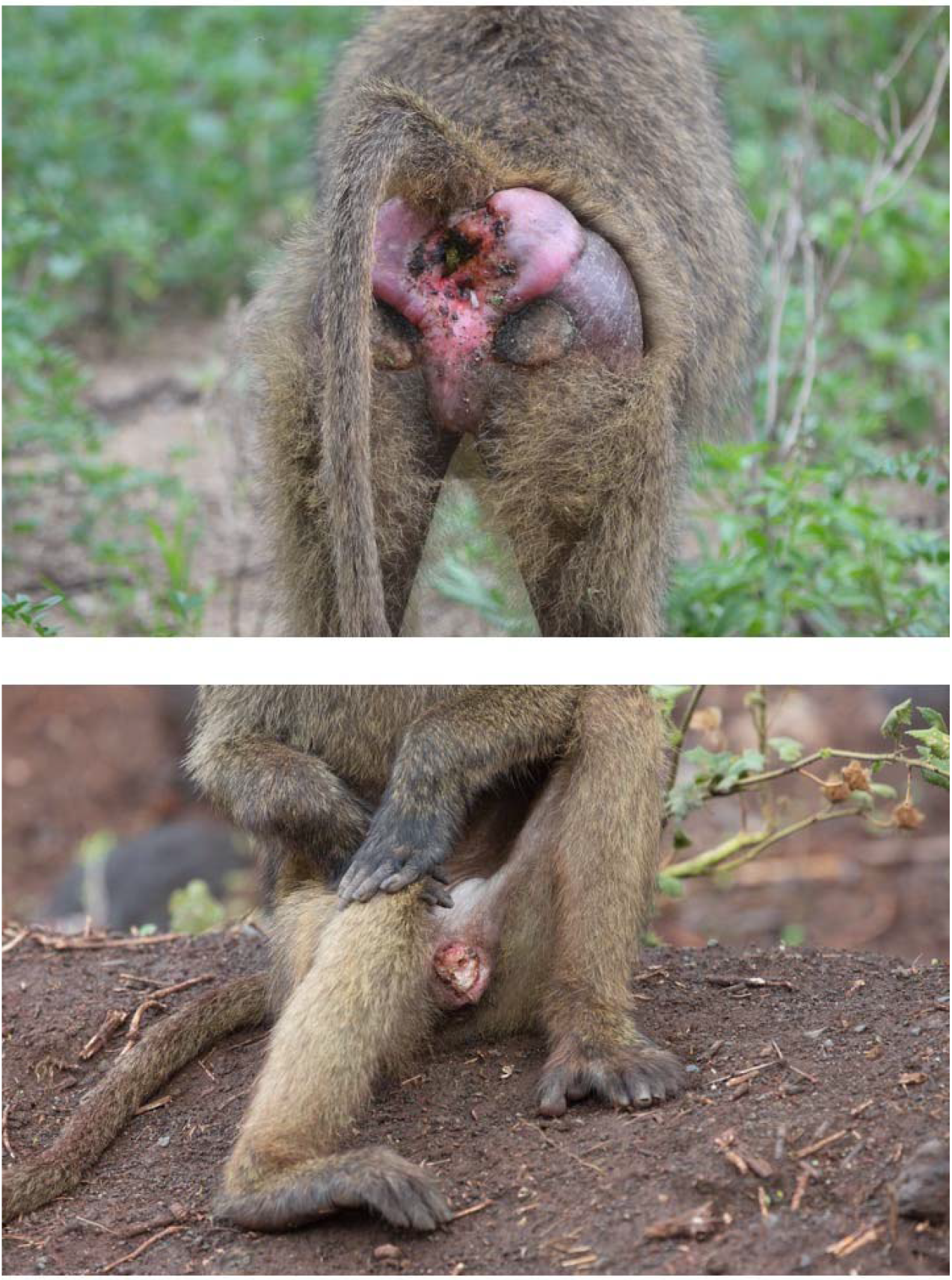
Genital skin ulcerations caused by *Treponema pallidum* subsp. *pertenue* in an adult female (top) and a subadult male (bottom) olive baboon at Lake Manyara National Park, Tanzania.

## 2 MATERIALS AND METHODS

This research adhered to the rules and regulations of the Tanzanian and German laws. The Animal Welfare and Ethics Committee of the German Primate Center approved the entire study.

### 2.1 Study site and subjects

Fieldwork was conducted at LMNP, Northern Tanzania, during two field seasons (April to December) in 2015 and 2016. Our baboon group was habituated during four months before the data collection phase of the study to facilitate behavioral observations from a distance of fewer than five meters.

The group consisted of approximately 170 individuals, of which 53 were adult and subadult females, 35 adult and subadult males and more than 70 immature individuals. In our analyses, we included 32 cycling females and their 29 male partners which could all be individually identified. The genital health status (GHS) was visually assessed and all adult and subadult individuals were classified as either genitally “ulcerated” or “non-ulcerated” using macroscopic visual cues (Knauf et al., 2012). Genital ulcerations could range from small-medium ulcers to a severe mutilation of the outer genitalia (Figure 1).

### 2.2 Behavioral data

We conducted full-day focal follows (Altmann, 1974) from dawn to dusk on 32 cycling females. To maximize the number of observed mating events, we focused on females in their peak estrus, denoted by maximal tumescence and bright pink color of their anogenital skin (Zinner et al., 2004). We collected 597 hours of observation data, with an average of 16.40 ± 10.02 hours (mean ± SD, range 1.50 – 39.00 hours) per focal female. We collected data on the number of mating events, type of copulation, and the presence of copulation calls, darting behavior and PCG (Table 1). Behavioral data were recorded in the field on a hand-held Samsung Galaxy using Pendragon 5.1.2 software (Pendragon Software Corporation, USA).

**TABLE 1.** Definition of variables

|  | Definition |
| --- | --- |
| **Copulation/mating event** | male mounting an estrous female and performing pelvic thrusts (with intromission*) and with or without ejaculation |
| **Type of copulation** | ejaculatory or non-ejaculatory: indicated by visible fresh sperm on the male's penis or by the sperm plug on the female's genitalia after copulation |
| **Pelvic thrusts** | number of male pelvic thrusts during copulation |
| **Copulation call** | context-specific calls female utter during, or at the end of a copulation |
| **Darting** | rapid withdraw in which a female can run away several meters from the male after copulation |
| **Post-copulatory grooming (PCG)** | male/female grooms the mating partner 15 seconds after copulation occurred |

\*Except for severely ulcerated males that lack the corpus penis as intromission cannot occur

### 2.3 Statistical analysis

We run generalized linear mixed models (GLMM, Baayen, 2008) to examine the post-copulatory behavior of our baboon population. All models were performed in R v3.4.4 (R Core Team 2018) with the lme4 package v 1.1-15 (Bates et al., 2015) and collinearity of the variables was checked using the package car (Fox & Weisberg 2011). Maximum likelihood ratio tests were used to test the full model with fixed factors against the null model (Faraway, 2006). Since interactions between the fixed predictors did not significantly improve any model fit, we excluded them from all models for parsimony and a more reliable interpretation of the main effects. In all models, female, male and pair identities were included as random factors.

### 2.5 Model description

#### Model 1: Copulation calls

The first model analyzed whether the occurrence of copulation calls was affected by the male or female GHS, the type of copulation, or the number of pelvic thrusts. The response variable was the presence or absence of copulation calls per mating event (1/0) with a binomial error structure and a logit link function.

#### Model 2: Darting behavior

With the second model, we examined whether post-copulatory darting was affected by the male or female GHS, type of copulation and the occurrence of copulation calls. The response variable was the presence or absence of darting per mating event (1/0) with a binomial error structure and a logit link function.

#### Model 3: Occurrence of post-copulatory grooming (PCG)

With the third model we investigated whether the occurrence of PCG is affected by the GHS of the male and female, respectively, presence of copulation calls and the type of copulation. Here the response variable was the presence or absence of PCG per mating event (1/0) with a binomial error structure and a logit link function.

#### Model 4: Duration post-copulatory grooming (PCG)

With this model, we examined whether the duration of PCG (in seconds) was affected by the presence of copulation calls and the type of copulation. Here we generated two GLMMs; one model for PCG performed by males (PCG-M) and another for PCG performed by females (PCG-F). Each model assumed that the duration of PCG depended on the presence of copulation calls and type of copulation. Both models were fitted using the glmmADMB package (Fournier et al., 2012) with a negative binomial error structure and a logit link function.

## 3 RESULTS

The prevalence of ‘genital ulcerated’ individuals in our study group (determined visually) remained relatively stable throughout the 18-months study period. Only three adult females and three adult males switched from ‘non-ulcerated’ to ‘ulcerated’ between field seasons. Therefore, at the end of the study, 44% (N=23) of the 53 adult and subadult females and 47% (N=17) of the 35 adult and subadult males displayed genital ulcers (Figure 1). Genital ulcers were observed in 40% (N=32) of the females participating in sexual interactions and in 53% (N=35) of the males. In total, we have observed 517 copulations among 32 females and 29 males. Evidence for ejaculation was found in 31.5% (N=163) of the copulations. Females uttered copulation calls in 25.5% (N=132), darting occurred in 41.7% (N=216) and PCG in 27.2% (N=141). The frequency of copulations calls and darting differed slightly between ejaculatory to non-ejaculatory copulations (Figure 2). But the likelihood of uttering copulation calls increased with ejaculatory mating (p<0.001, Table 2). Males who lacked the corpus penis were observed ejaculating towards the ground or against their legs as there was no way to direct the sperm into the female’s genital tract. Nevertheless, two females in our study group uttered copulation calls even when mating with males lacking the corpus penis, where intromission was not observed (i.e., males solely performed pelvic thrusts).

**FIGURE 2.**
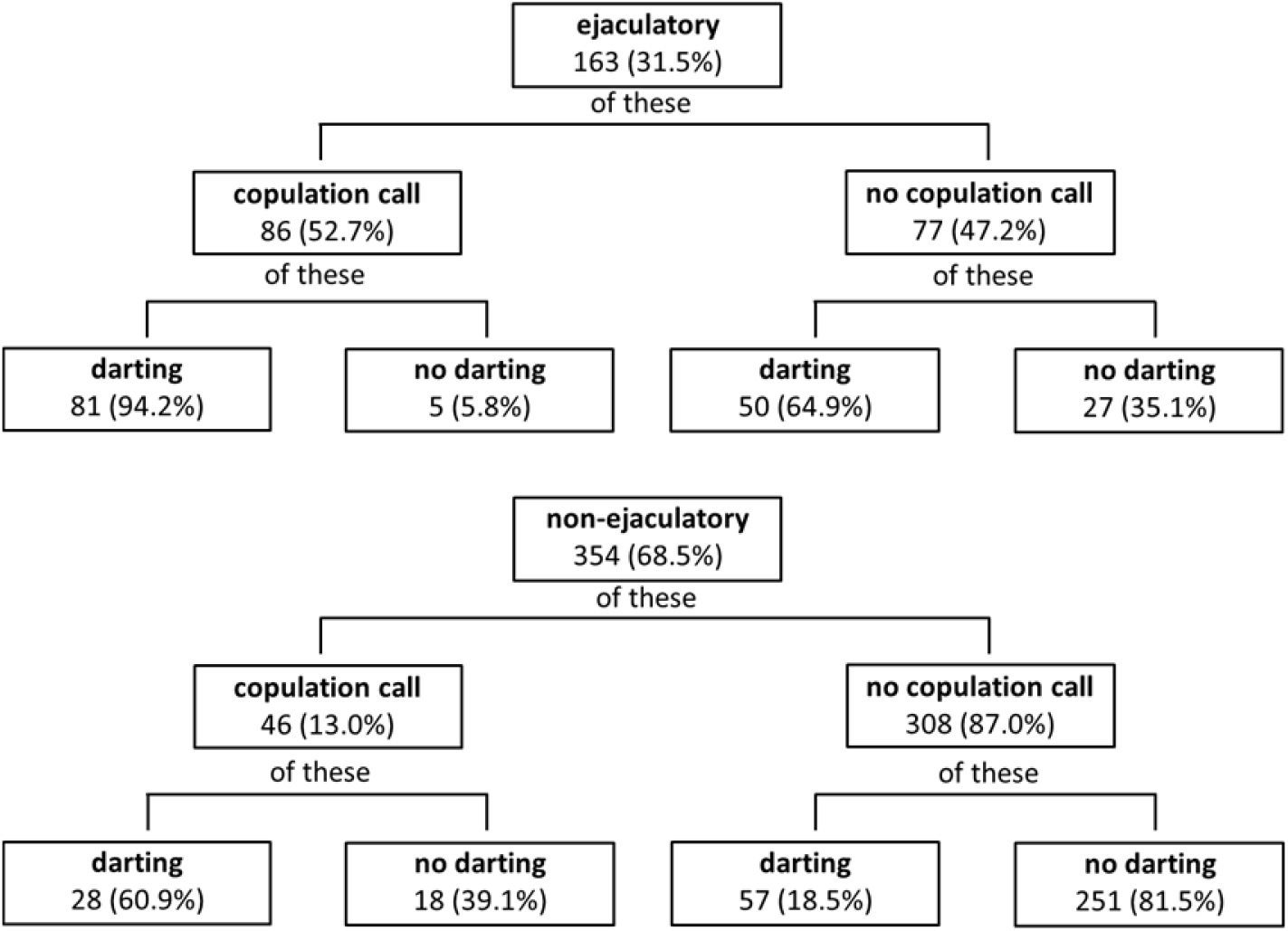
Frequency of copulation calls and darting after ejaculatory copulations (top; n = 163) and non-ejaculatory copulations (bottom; n = 354). Total number of copulations = 517.

**TABLE 2.** Copulation call model. Binary GLMM evaluating if the likelihood of uttering a copulation call is affected by the male and female genital health status (GHS), the number of pelvic thrusts and the type of copulation. Estimates, standard errors (SE), z-values, and 2.5% and 97.5% confidence intervals (CI) are shown for fixed effects. Intercept with a reference category for non-ulcerated individuals and non-ejaculatory events.

| Term | Estimate | SE | CI lower | CI upper | z value | P |
| --- | --- | --- | --- | --- | --- | --- |
| Intercept | -3.725 | 0.830 | -5.639 | -2.245 | -4.490 | - |
| Male GHS | 0.057 | 0.803 | -1.518 | 1.758 | 0.071 | 0.944 |
| Female GHS | 1.482 | 0.945 | -0.347 | 3.553 | 1.569 | 0.117 |
| Type of copulation | 2.800 | 0.430 | 2.005 | 3.706 | 5.759 | <0.001 |

Darting was observed in 80% (N=131) of the copulations with ejaculation, in contrast to only 24% (N=85) of the non-ejaculatory copulations. Darting was also more frequent when females uttered copulation calls (94%, N= 81). These observations were corroborated by our second model. The likelihood of darting was higher when females gave copulation calls and when the male ejaculated (p<0.001, Table 3). Darting never led to the termination of a consortship as males kept track of the females, even if the female covered distances of more than 10 meters. Moreover, consort take-overs were rarely observed in our group (n=7 over the 18-months study period).

**TABLE 3.** Post-copulatory darting model. Binomial GLMM evaluating if the likelihood of darting is influenced by the male and female genital health status (GHS), presence of copulation calls and type of copulation. Estimates, standard errors (SE), z-values, and 2.5% and 97.5% confidence intervals (CI) are shown for fixed effects. Intercept with reference category for non-ulcerated individuals, absence of copulation calls and non-ejaculatory events.

| Term | Estimate | SE | CI lower | CI upper | z value | P |
| --- | --- | --- | --- | --- | --- | --- |
| Intercept | -1.468 | 0.503 | -2.537 | -0.497 | -2.919 | - |
| Male GHS | -0.525 | 0.621 | -1.714 | 0.770 | -0.844 | 0.118 |
| Female GHS | 0.883 | 0.565 | -0.242 | 2.092 | 1.563 | 0.398 |
| Copulation call | 2.408 | 0.480 | 1.502 | 3.402 | 5.017 | <0.001 |
| Type of copulation | 2.500 | 0.380 | 1.781 | 3.277 | 6.586 | <0.001 |

In most cases (72.8%, N= 376), no PCG occurred. When it occurred, males initiated PCG more often than females regardless of the type of copulation (Table 4). The occurrence of PCG was neither affected by the occurrence of copulation calls nor by the type of copulation (Table 5). However, the duration of PCG performed by males was longer when females uttered copulation calls (p=0.019). No effect was found for PCG initiated by females (Table 6).

**TABLE 4.** Frequency of post-copulatory grooming (PCG) initiation in relation to copulation type (number of cases in parentheses).

|  | Copulations<br>(512)* | Ejaculatory<br>30.6 % (159) | Non-ejaculatory<br>68.2 (353) |
| --- | --- | --- | --- |
| **No PCG** | 72.7% (376) | 77.3 % (123) | 71.46% (253) |
| **Male initiated** | 17.7% (92) | 15 % (24) | 19.2 % (68) |
| **Female initiated** | 8.5% (44) | 4.5 % (12) | 9 % (32) |

\*Five copulations excluded from the total (n= 517) as PCG could not be assessed properly

**TABLE 5.**
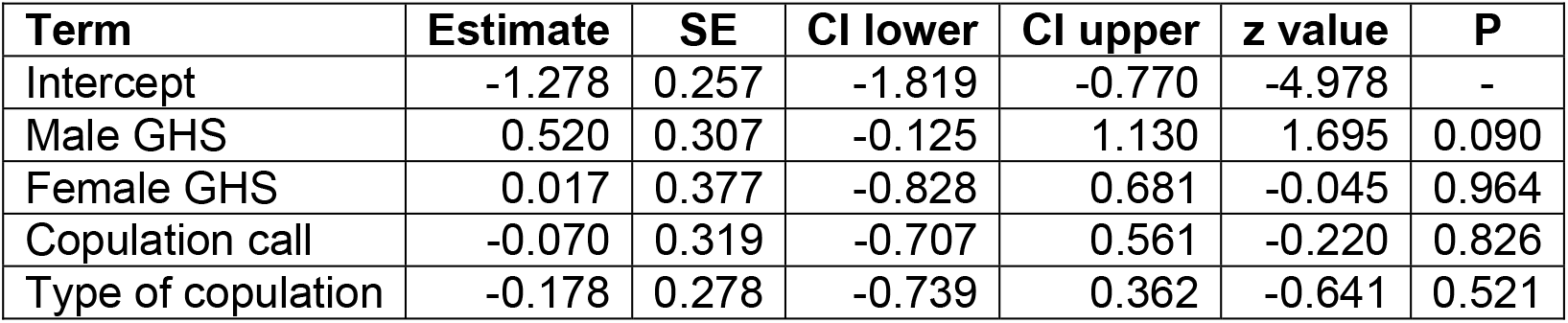
Post-copulatory grooming (PCG)-presence model. Binomial GLMM evaluating if the likelihood of PCG is affected by the male and female genital health status (GHS), presence of copulation calls and type of copulation. Estimates, standard errors (SE), z-values, and 2.5% and 97.5% confidence intervals (CI) are shown for fixed effects. Intercept with reference category for non-ulcerated individuals, absence of copulation calls and non-ejaculatory events.

| Term | Estimate | SE | CI lower | CI upper | z value | P |
| --- | --- | --- | --- | --- | --- | --- |
| Intercept | -1.278 | 0.257 | -1.819 | -0.770 | -4.978 | - |
| Male GHS | 0.520 | 0.307 | -0.125 | 1.130 | 1.695 | 0.090 |
| Female GHS | 0.017 | 0.377 | -0.828 | 0.681 | -0.045 | 0.964 |
| Copulation call | -0.070 | 0.319 | -0.707 | 0.561 | -0.220 | 0.826 |
| Type of copulation | -0.178 | 0.278 | -0.739 | 0.362 | -0.641 | 0.521 |

**TABLE 6.** PCG duration model. GLMMs evaluating if the duration of PCG is affected by the presence of copulation calls and type of copulation. Estimates, standard errors (SE), z-values, and 2.5% and 97.5% confidence intervals (CI) are shown for fixed effects. PCG performed by males and females is shown in PCG-M and PCG-F respectively. Intercept with reference category for non-ulcerated individuals, absence of copulation calls and non-ejaculatory events (GHS = genital health status).

| Term | Estimate | SE | CI lower | CI upper | z value | P |
| --- | --- | --- | --- | --- | --- | --- |
| **PCG-M** |  |  |  |  |  |  |
| Intercept | 5.213 | 0.185 | 4.851 | 5.575 | 28.24 | - |
| Male GHS | 0.106 | 0.191 | -0.268 | 0.480 | 0.555 | 0.579 |
| Female GHS | -0.197 | 0.180 | -0.550 | 0.156 | -1.091 | 0.275 |
| Copulation call | 0.472 | 0.201 | 0.078 | 0.864 | 2.352 | **0.019** |
| Type of copulation | 0.029 | 0.198 | -0.359 | 0.417 | 0.147 | 0.883 |
| **PCG-F** |  |  |  |  |  |  |
| Intercept | 5.229 | 0.229 | 4.780 | 5.677 | 22.84 | - |
| Male GHS | 0.337 | 0.327 | -0.304 | 0.978 | 1.029 | 0.304 |
| Female GHS | -0.124 | 0.331 | -0.773 | 0.525 | -0.375 | 0.708 |
| Copulation call | -0.147 | 0.456 | -0.485 | 0.780 | 0.456 | 0.648 |
| Type of copulation | 0.164 | 0.588 | -0.381 | 0.709 | 0.588 | 0.556 |

## 4 DISCUSSION

The impact of sexually transmitted infections on the mating behavior of nonhuman primates is still poorly understood. Our data from the LMNP baboons suggest that genital ulcers have an impact on female mate choice and male mating performance (Paciência et al., 2019), whereas the post-copulatory behavior seems to remain unaffected by the presence of genital ulcers.

Female olive baboons at LMNP produce copulation calls less often than other baboon species (yellow baboons: 80%, Collins, 1981; 96.9%, Semple, 1998; chacma baboons: 83%, Saayman, 1970; O’Connell & Cowlishaw 1994; Guinea baboons: 39%, Boese, 1973; olive baboons: 19%, Ransom, 1981; 62%, Bercovitch. 1985, 25% this study; but not hamadryas baboons: 18%, Swedell & Saunders 2006; 26.1% Nitsch et al., 2011). Moreover, female olive baboons at LMNP uttered copulation calls more likely when mating was followed by ejaculation. Similar findings were observed in previous studies, where copulation calls occurred more frequently, or had a longer duration after ejaculatory copulations (Saayman, 1970; Deputte & Goustard 1980; Collins, 1981; Todt et al., 1995; O’Connell & Cowlishaw 1994; Walz, 2016, but see Semple et al., 2002). Copulation calls have been suggested to constitute a mechanism to incite male-male competition in chacma baboons (O’Connell & Cowlishaw 1994; Crockford et al., 2007). In olive baboons, however, due to the long-term consortships (i.e. during the estrous periods) and even “friendships” (i.e., outside the estrus period (Smuts, 1985)), it was proposed that copulation calls function to reassure consortship formation and/or continuation with mating partners (Walz, 2016). According to the female choice hypothesis, calls can be directed to the current partner to encourage mate-guarding or to continue copulating (Todt et al., 1995). This can lead to a reduction of the likelihood of copulating with other partners and an increase in paternity certainty in males preferred by the female (Maestripieri & Roney 2005).

The occurrence and frequency of darting behavior is highly variable among olive baboons (25%, Ransom, 1981; 92%, Bercovitch, 1985; 76%, Walz, 2016, 41.7%, this study) as well as in chacma baboons (78%, Hall, 1962; 75%, Hall & DeVore 1965; 86–89%, Saayman, 1970; 78%, O’Connell & Cowlishaw 1995). In our study group, females darted more often after an ejaculatory mating and when the female uttered copulation calls. Similar observations were reported in another population of olive baboons, where females darted longer distances after ejaculatory copulations (Walz, 2016). While darting distance in chacma baboons was not affected by the occurrence of ejaculation, it was positively correlated with the duration of copulation calls (O’Connell & Cowlishaw 1995). In olive baboons, females have been described to run immediately after the copulation from the mating male towards another male (Hall & DeVore 1965), leading to consort take-overs (Smuts, 1985). Such behavior was not observed at LMNP as the darting female was usually followed by the consorting male, and consort take-overs were seldom observed. This might be due to the long term bonds observed between males and females in estrus, as females would frequently mate with the same male during different cycling periods (Paciência et al., 2019).

The function of post-copulatory grooming (PCG) is still unclear. Males might use PCG to prevent females from mating with other males and reduce sperm competition (Berenstain & Wade 1983; Kuester & Paul 1992; Nurnberg et al., 1994; Sonnweber et al., 2015). On the other hand, females might employ PCG as a means to either stimulate or avoid mating with the same male (Slob et al., 1986; Bancroft, 2005; Gumert, 2007) or decrease the risk of harassment by males (Smuts, 1985). Quantitative studies on PCG in baboons are scarce. In olive baboons, PCG presence has been related to the quality of the social bonds between the mating partners (Smuts, 1985). Yet, in chacma baboons, it is more frequently performed by females than males, except for females in their “swollen phase”, (i.e. maximum turgescent phase), where the grooming frequency by the male partners is higher (Saayman, 1970). In Barbary macaques, where PCG has been studied extensively, this behavior is known to occur in half of the mating events (Taub, 1980; Small 1990; Kuester & Paul 1992; Sonnweber et al., 2015). In this species, males are more likely to groom females after ejaculatory copulations, while females groom more often males after non-ejaculatory events (Sonnweber et al., 2015). This stands in contrast to our data from olive baboons at LMNP, where PCG did not take place in the majority of the cases and was neither affected by the type of copulation nor the presence of copulation calls. However, PCG duration performed by males was affected by female copulation calls irrespective of the type of copulation. Because copulation calls give hints to all males in the group that copulation just occurred, it might lead to male-male competition, and thus, males might be willing to increase their grooming bouts to prevent females from searching subsequent copulations with other males, which could aid in reducing sperm competition.

The clear comprehension of the possible functions of post-copulatory behaviors in nonhuman primates is still missing. In order to fill this gap, it is essential to collect and share quantitative data on sexual behavior and mating patterns of nonhuman primates. This is particularly important when group living species are affected by a sexually transmitted infection that has an impact on both sociality and reproduction leading to altered group dynamics.

## ACKNOWLEDGMENTS

We thank Tanzania Wildlife Research Institute (TAWIRI), especially J. D. Keyyu and R. Fyumagwa; Tanzania National Parks (TANAPA), especially I. A. V. Lejora and Y. Kiwango; LMNP headquarters staff, particularly R. Kaitila. A special thanks to the LMNP rangers who helped with fieldwork: P. Mkama, D. Baluya, P. Mbaryo and J. Bitulo. We also thank the Tanzania Commission for Science and Technology (COSTECH) for their support. Financial support was provided by the German Science Foundation (DFG), and conducted as part of the research group Sociality and Health in Primates (KN1097/4-1 to SK and ZI548/5-1 to DZ).

## AUTHOR CONTRIBUTIONS

F.M.D.P, S.K and D.Z designed the study. F.M.D.P, I.S.C, I.F.L and S.K performed field work. F.M.D.P collected the data in the field. Data analysis was done by F.M.D.P. The paper was written by F.M.D.P, S.K and D.Z.

## DATA AVAILABILITY STATEMENT

The data that support the findings of this study are available from the corresponding author upon reasonable request.

